# Early transcriptional divergence underlies cell fate bias in bovine embryos

**DOI:** 10.1101/2025.07.01.661069

**Authors:** Koyama Hinata, Daisuke Mashiko, Masahiro Kaneda, Atchalalt Khurchabilig, Satoshi Sugimura

## Abstract

Developmental plasticity, or the ability of early embryonic cells to contribute to multiple lineages, is traditionally considered equal among sister blastomeres during early cleavage. However, divergence may occur earlier than expected. We performed single-cell RNA sequencing of bovine embryos from the 2- to 8-cell stages to examine transcriptional asymmetry. While gene expression was uniform at the 2-cell stage, variability increased at the 4-cell stage and became pronounced by the 8-cell stage. At this stage, blastomeres showed heterogeneity in MAPK pathway genes (e.g., *RAC1*, *MAPK14*) and the trophectoderm marker *CDX2*. These differences were associated with blastomere size; larger blastomeres more frequently initiated cavity formation, a functional marker of trophectoderm fate. Our findings suggest that both molecular and physical asymmetries contribute to early lineage bias, and that developmental plasticity may be lost in an asynchronous, cell-specific manner before visible morphological events such as compaction.

## Introduction

Early mammalian embryos exhibit a high degree of developmental plasticity, whereby individual blastomeres retain the capacity to generate both embryonic and extra-embryonic lineages. This flexibility reflects the transient totipotent state during the early cleavage stages, in which each blastomere can theoretically give rise to a complete organism.

This plasticity has traditionally been considered equivalent between sister cells during early cleavage stages^1, 2^ with this property maintained until the morula stage, when compaction and positional cues trigger the initial lineage segregation^3, 4^. However, accumulating evidence suggests that developmental potential is restricted earlier than previously assumed, with molecular asymmetries and lineage biases arising during the early cleavage stages. Developmental plasticity is tightly regulated by pluripotency networks and extrinsic signals, such as the fibroblast growth factor and Wnt pathways^5^. Therefore, symmetry-breaking processes are possibly initiated well before apparent morphological distinctions.

In mice, developmental bias has been reported as early as the 2-cell stage, with one blastomere contributing more to the inner cell mass (ICM) and the other favouring the trophectodermal fate^6, 7^. These early tendencies are accompanied by differences in gene expression, cell cycle dynamics, and epigenetic states, even before overt polarity and spatial organization^8, 9^. Single-cell RNA-sequencing (scRNA-seq) studies revealed that transcriptional asymmetries between sister blastomeres emerge during cleavage in mouse embryos. These asymmetries, reported as early as the 2-cell stage in some studies, are associated with early cell fate bias, possibly representing the earliest molecular signatures of lineage segregation^8, 10, 11^. Although the robustness and reproducibility of such early transcriptomic differences have been questioned^12^, these reports suggest that symmetry breaking at the molecular level precedes and influences the initial cell fate decisions in mammals.

Whether a similar early divergence occurs in non-rodent mammals developmentally analogous to humans remains unclear. Bovine embryos offer a biologically relevant model to evaluate this, as they closely resemble human embryos in terms of cleavage dynamics, timing of zygotic genome activation, and gradual segregation of embryonic and extraembryonic lineages^13–17^. These features make cattle valuable complementary models to rodents for early cell fate studies.

To date, most studies have analysed inter-blastomere transcriptional variation at a single time point or within pooled embryos, limiting our understanding of the ways in which asymmetries evolve across developmental stages in individual embryos. Only a few studies have attempted to track the transcriptional heterogeneity between blastomeres in a stage-resolved embryo-specific manner^10, 11^ and even fewer have extended this analysis beyond the early cleavage stages.

To address the above-mentioned limitations, we performed scRNA-seq of the individual blastomeres of bovine embryos at the zygote, 2-cell, 4-cell, and 8-cell stages. By capturing cell-specific transcriptomic profiles across successive cleavage divisions, we aimed to determine when the transcriptional divergence between sister blastomeres first emerges and whether these differences correspond to the onset of lineage bias. We found that divergence begins at the 4-cell stage and becomes pronounced at the 8-cell stage, coinciding with early signs of lineage bias. Our findings provide new insights into the progressive restriction of developmental plasticity before the morula stage in a non-rodent mammalian model, enhancing our understanding of the timing of and mechanisms underlying early cell fate decisions.

## Results

### Progressive divergence in gene expression between sister blastomeres from the 4-cell stage

We successfully generated SMART-seq libraries from 90 individual blastomeres derived from ten zygotes (1-cell), ten 2-cell, five 4-cell, and five 8-cell stage embryos. Each library set represents a complete group of sister blastomeres from the same embryo, allowing precise comparisons of gene expression variability within and between embryos across developmental stages (Figure 1a). We performed principal component analysis (PCA) and hierarchical clustering to assess the transcriptomic variability across developmental stages. Blastomeres of different stages (1-, 2-, 4-, and 8-cell stages) formed distinct stage-specific clusters (Figure 1b). At the 2-cell stage, gene expression profiles were highly similar between embryos and sister blastomeres, indicating minimal variability. However, transcriptomic divergence increased as the development progressed. By the 4-cell stage, variability was observed both within and between embryos, which became more pronounced at the 8-cell stage.

**Figure 1.**
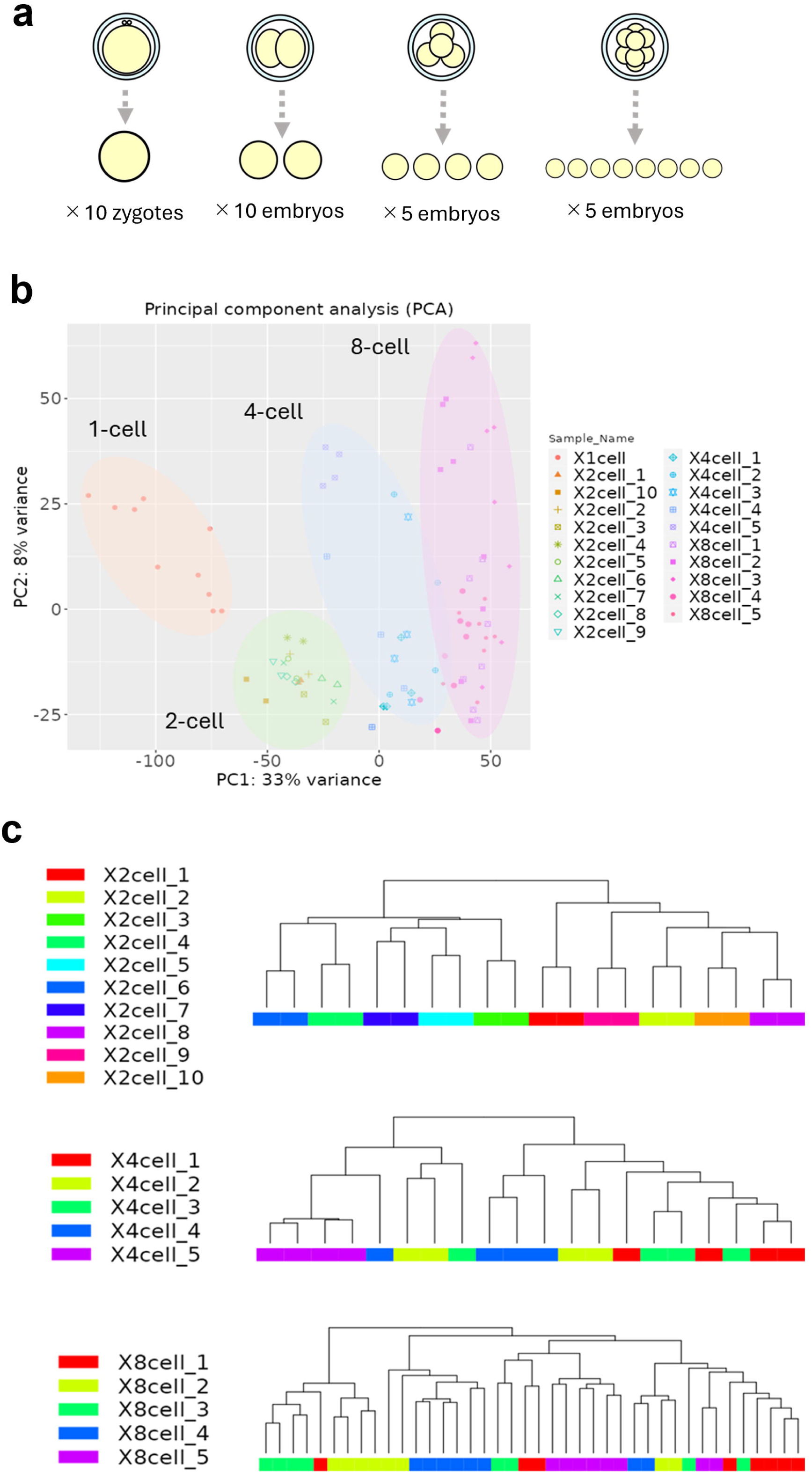
Transcriptomic profiling of individual blastomeres across early developmental stages. (a) Schematic showing the isolation of individual blastomeres from bovine embryos at the 1-cell (n=10), 2-cell (n=10), 4-cell (n=5), and 8-cell (n=5) stages. All blastomeres from each embryo were collected (total 90 cells), enabling within- and between-embryo comparisons using SMART-seq. (b) Principal component analysis (PCA) of individual blastomeres isolated from the 1-, 2-, 4-, and 8-cell stage embryos. Blastomeres of different developmental stages are plotted according to the first two principal components. (c) Hierarchical clustering of individual blastomeres based on the global gene expression profiles. Each branch represents an individual blastomere, with blastomeres derived from the same embryo indicated by the same colour.

Cluster analysis supported these observations (Figure 1c). Although sister blastomeres clustered tightly at the 2-cell stage, such clustering was less pronounced at the 4-cell stage. By the 8-cell stage, blastomeres of the same embryo did not cluster together. Instead, each blastomere formed a separate branch, indicating substantial intercellular heterogeneity. These results suggest that transcriptional variability is initially low but increases with each cleavage division, supporting the notion that molecular divergence begins as early as the 4-cell stage.

### MAPK and related signalling pathway enrichment of genes showing expression variability at the 8-cell stage

Next, we used the ratio of within-embryo to total gene expression variance (SSwe/SS; see Methods, *Assessment of transcriptional asymmetry*) to quantify transcriptional asymmetry between sister blastomeres. As illustrated in Figure 2a, this approach partitions total variance into variability within embryos (SSwe) and between embryos (SSbe). The top panel represents high SSwe with low SSbe, indicating substantial heterogeneity between blastomeres within the same embryo. In contrast, the bottom panel shows low SSwe and high SSbe, reflecting uniform expression within embryos but greater differences between embryos (Figure 2a). At the 2-cell stage, distribution of SSwe/SS values exhibited a bimodal pattern, with peaks observed near 0 and 1, indicating the coexistence of symmetric and highly asymmetric gene expression between blastomeres. As development progressed to the 4- and 8-cell stages, this distribution shifted toward intermediate SSwe/SS values, suggesting a broader and more systematic increase in transcriptional divergence. By the 8-cell stage, several genes exhibited moderate-to-high inter-blastomere variability (Figure 2b).

**Figure 2.**
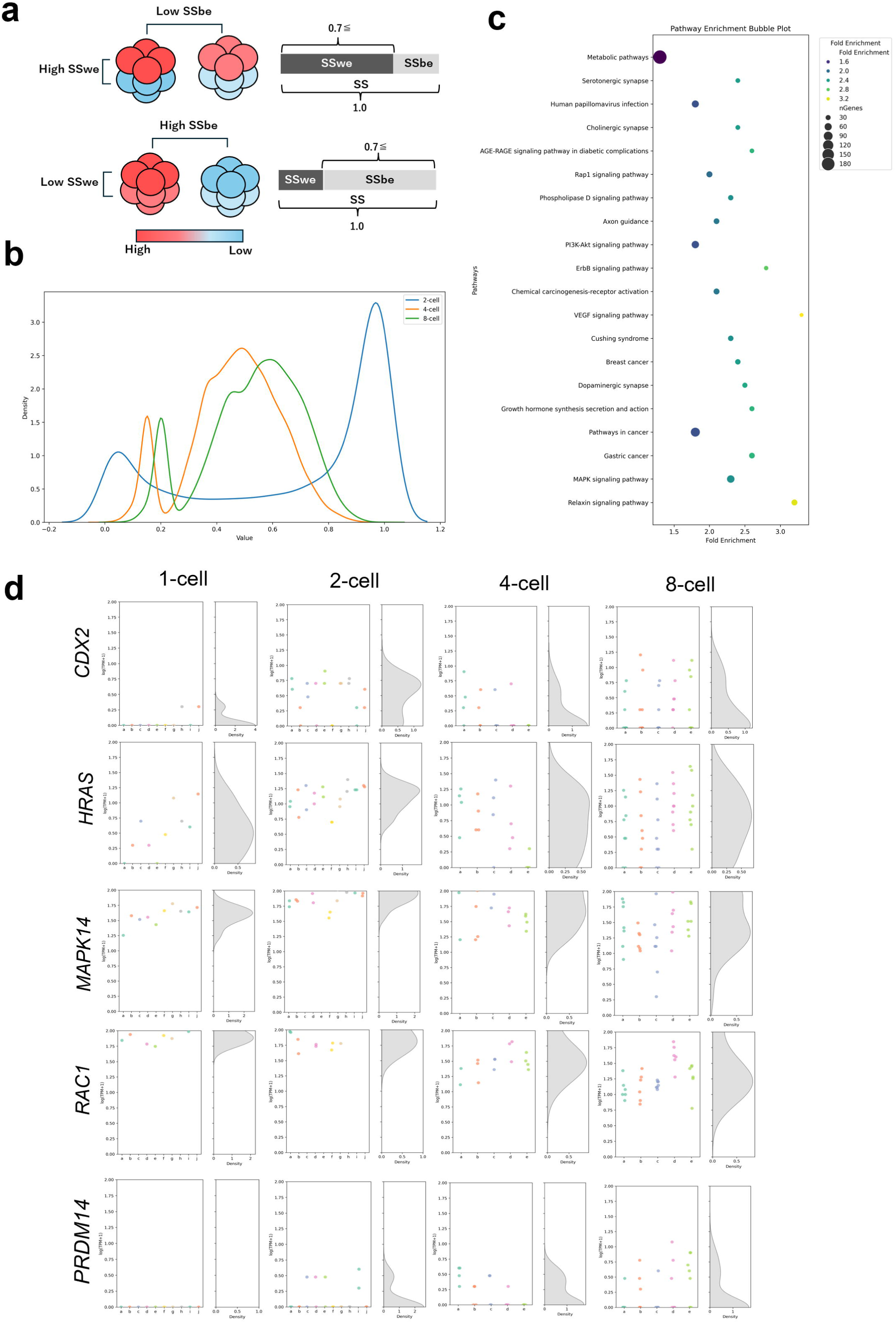
Assessment of the transcriptional asymmetry between sister blastomeres. (a) Conceptual illustration of gene expression variance (total sum of squares, SS) partitioned into within-embryo (SSwe) and between-embryo (SSbe) components for genes with high SSwe/SS ratios (≥ 0.70 at the 8-cell stage). The upper panel illustrates high SSwe with low SSbe, while the lower panel shows low SSwe with high SSbe. Colors indicate relative expression levels. (b) Distribution of SSwe / SS values of genes at the 2-, 4-, and 8-cell stages. SS represents the total variance in gene expression across all blastomeres, whereas SSwe represents the variance specifically between sister blastomeres within the same embryo. SSwe/SS ratio indicates the proportion of total variance attributable to within-embryo variability, with values near 0 indicating symmetric expression, and those near 1 indicating asymmetric expression between sister blastomeres. (c) KEGG pathway enrichment analysis of genes exhibiting the highest SSwe/SS values at the 8-cell stage. (d) Comparison of the expression levels of the representative early lineage specification-associated genes (*CDX2*, *HRAS*, *MAPK14*, *RAC1* and *PRDM14*) between sister blastomeres.

We identified the genes with the highest SSwe/SS values at the 8-cell stage (Supplementary Data 1) and performed KEGG pathway enrichment analysis. The identified genes were significantly enriched in the MAPK and related pathways, including VEGF, ErbB, AGE-RAGE, Rap1, and PI3K-Akt signaling pathways (Figure 2c). Notably, these pathways were not enriched in blastomeres at the 2- and 4-cell stages (Supplementary Data 1 and Supplementary Figure 1).

Several genes with high SSwe/SS ratios were associated with known regulators or markers of lineage specification, including the trophectoderm (TE)-associated genes, *RAC1* and *CDX2* ^18^, and *HRAS* and *MAPK14*, which are implicated in the MAPK-dependent regulation of CDX2 expression^19^. PRDM14, a gene linked to ICM specification in mouse embryos^20^, also exhibited stage-dependent expression divergence.

We compared the expression levels of these genes between sister blastomeres (Figure 2d). *CDX2* expression was first detected at the 2-cell stage, with inter-blastomere variability emerging at the 4-cell stage and persisting through the 8-cell stage. *HRAS*, *MAPK14*, and *RAC1* were already expressed from 1-cell and showed relatively uniformly expression at the 2-cell stage; however, expression differences between sister blastomeres became more pronounced from the 4-cell to the 8-cell stage. *PRDM14* transcripts were also detected from the 2-cell stage, with inter-blastomere variability gradually increasing from the 2-cell to the 8-cell stage.

To further explore the relationship between MAPK signalling and *CDX2* expression, we stratified the 8-cell blastomeres into high and low CDX2-expressing groups and performed PCA. High CDX2-expressing were clustered tightly along the first principal component (PC1; Figure 3a). KEGG analysis of the differentially expressed genes (DEGs) between the two groups showed enrichment of MAPK and its associated signalling pathways (Supplementary Data 2 and Figure 3b). Positive correlations were observed between *CDX2* and *RAC1*, *HRAS*, and *MAPK14* levels (Figure 3c). These findings suggest that the transcriptional divergence between sister blastomeres becomes evident at the 4-cell stage, increases at the 8-cell stage, and is accompanied by the differential expression of signalling pathway components, most notably those related to MAPK, which possibly contribute to early TE lineage specification.

**Figure 3.**
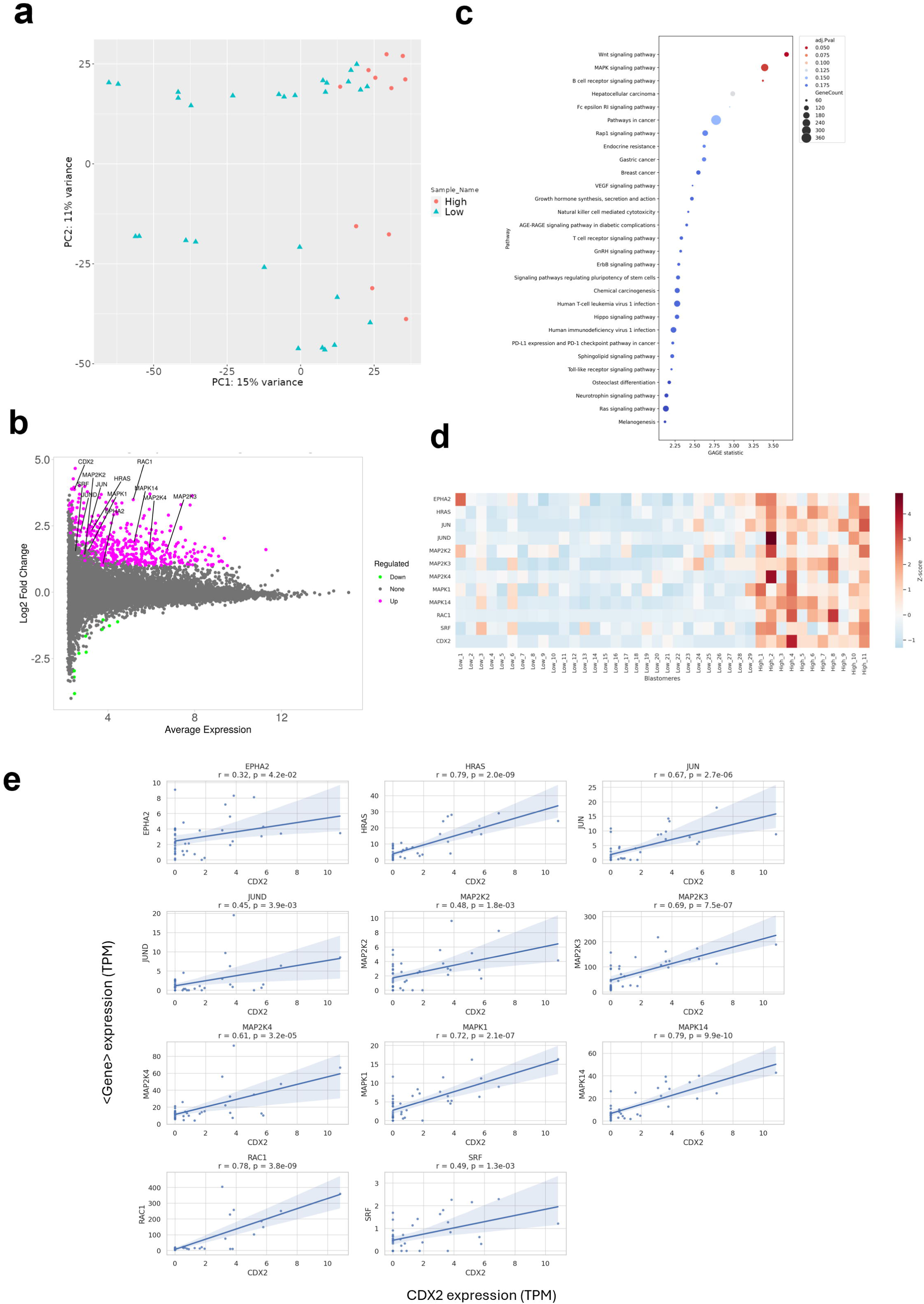
Comparative analysis of 8-cell stage blastomeres stratified by CDX2 expression levels. (a) PCA of 8-cell stage blastomeres classified into high and low CDX2-expressing groups. (b) KEGG pathway enrichment analysis of the differentially expressed genes between the high and low CDX2-expressing blastomeres. (c) Correlation analysis between CDX2 and *HRAS*, *MAPK14*, and *RAC1* levels in 8-cell stage blastomeres.

### Size-associated transcriptional differences and cavity formation

To determine whether blastomere size is associated with the molecular differences relevant to lineage specification, we performed differential gene expression analysis of the largest and smallest sister blastomeres at the 2-, 4-, and 8-cell stages. No DEGs were detected at the 2-cell stage; however, several DEGs emerged at the 4- and 8-cell stages (Supplementary Table 1). Notably, *CDX2* and *RAC1* levels were upregulated in the largest blastomeres at both stages, whereas *HRAS* and *MAPK14* levels were higher in the largest blastomeres than in the smallest blastomeres at the 8-cell stage (Figure 4b). *HRAS*, *MPAK14*, and *RAC1* were also highly expressed at the zygote stage (Figure 2c), suggesting a maternal origin. These results suggest that large blastomeres are transcriptionally biased toward the activation of TE-associated pathways, such as the MAPK pathway, indicating the potential unequal inheritance of maternal transcripts.

**Figure 4.**
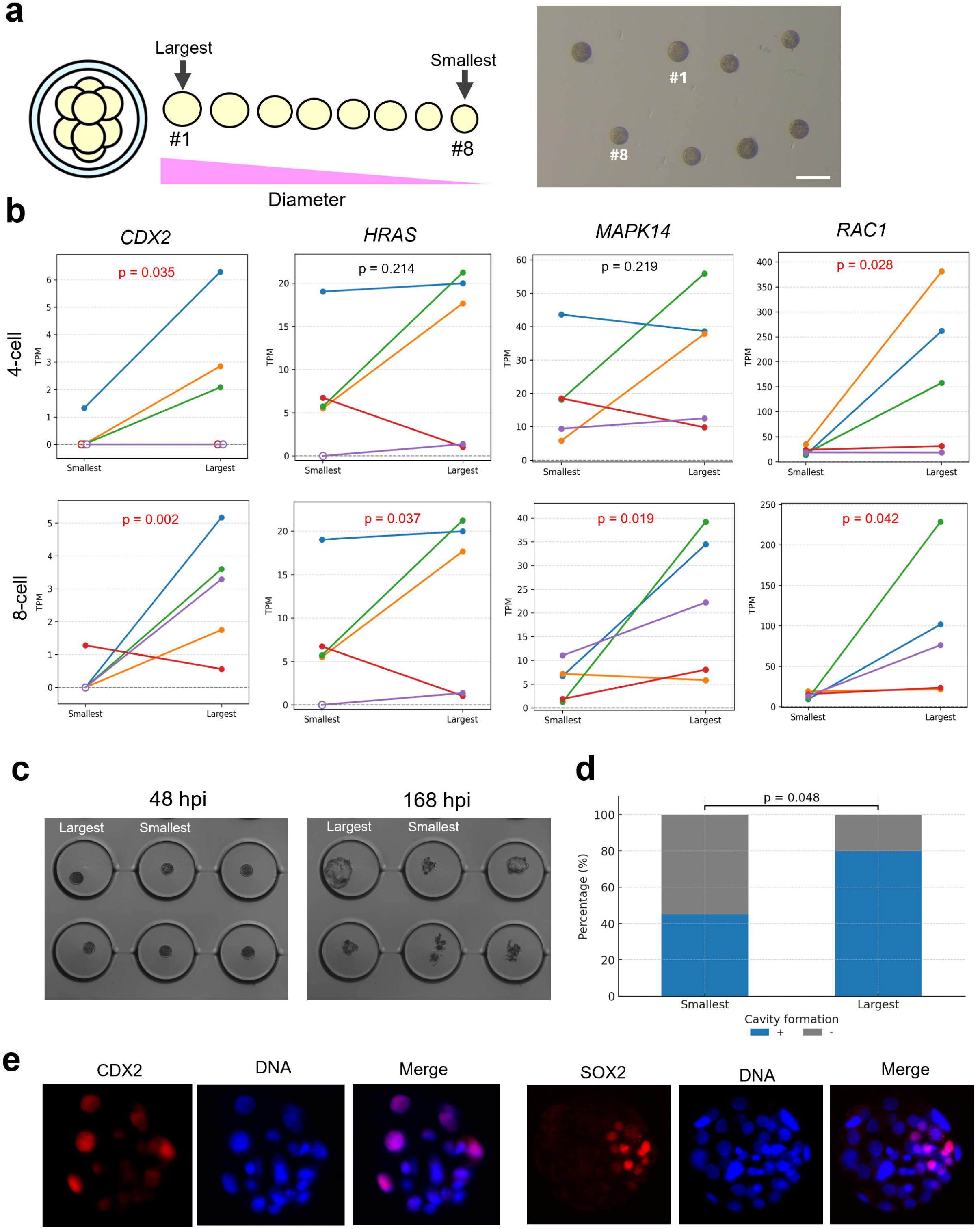
Size-associated differences in gene expression and cavity formation in blastomeres. (a) Representative images of individual blastomeres isolated from an 8-cell stage embryo. The blastomeres were numbered sequentially from left to right for identification. The largest blastomere was designated as #1, whereas the smallest was designated as #8. Scale bar, 100Lμm. (b) Differential expression analysis of *CDX2*, *HRAS*, *MAPK14*, and *RAC1* between the largest and smallest blastomeres at the 4- and 8-cell stages. Each colour represents a pair of largest and smallest blastomeres derived from the same embryo. Red P-values indicate the statistical significance. (c) Representative images of the blastomeres immediately after isolation at 48 h post-insemination (hpi) and at 168 hpi. Cavity formation in the well is indicated by a circle. (d) Quantification of the proportion of embryos that formed a cavity at any point during the 192 hpi period, in those derived from the largest and smallest blastomeres. (e) CDX2 and SOX2 expression levels in pseudo-blastocysts derived from the largest blastomere at 192 hpi.

Next, we examined whether this transcriptional bias corresponds to functional differences in developmental behaviour by assessing cavity formation, a morphological feature typically associated with TE activity, in size-classified blastomeres (Figure 4c and d). Cavity formation at any point during the 192 hours post-insemination (hpi) period was observed in 80% (16/20) of embryos derived from the largest blastomeres, compared to 45% (9/20) from the smallest, indicating a significant difference. At 192 hpi, cavity formation was maintained in 45% (9/20) of embryos derived from the largest blastomeres, which were subsequently analyzed by immunofluorescence staining. On average, these structures contained 15.3 CDX2-positive (TE) cells and 6 SOX2-positive (ICM) cells. In contrast, only 2 out of 20 embryos (10%) derived from the smallest blastomeres exhibited cavity formation at 192 hpi, and none were available for marker analysis. Therefore, the largest blastomeres not only initiated and maintained cavity formation but also underwent differentiation into TE lineages, as evidenced by CDX2 expression. Moreover, detection of SOX2-positive cells indicated that these blastomeres retained developmental plasticity, maintaining their ability to contribute to both the TE and ICM lineages. Collectively, these results suggest that blastomere size is linked to both the transcriptional and functional properties relevant to early lineage bias.

## Discussion

Early cleavage stage embryos are traditionally considered as collections of equivalent blastomeres, each retaining the capacity to equally contribute to the embryonic and extraembryonic lineages until the onset of compaction^3, 4^. However, studies on mouse embryos have challenged this view by suggesting that early cleavage divisions can generate asymmetries in gene expression, cell behaviour, and developmental potential^8, 10, 11, 21, 22^. However, whether a similar early divergence occurs in non-rodent mammals remains unclear.

Here, scRNA-seq analysis revealed that the transcriptional divergence between sister blastomeres was detectable at the 4-cell stage but intensified at the 8-cell stage in bovine embryos. Notably, MAPK signalling pathway-associated genes, including *RAC1*, *HRAS*, and *MAPK14*, exhibited substantial inter-blastomere variability at the 8-cell stage. Considering the role of MAPK signalling in regulating CDX2 expression and TE specification^19^, this early transcriptional asymmetry suggests the onset of lineage bias before morphological polarisation, such as compaction. Importantly, these MAPK-associated genes were already expressed at the zygote (1-cell) stage, prior to major embryonic genome activation (EGA), which occurs at the 8-cell stage in bovine embryos ^23^. This indicates that the observed variability likely originates from differential inheritance or depletion of maternal transcripts, rather than *de novo* transcription driven by EGA. Thus, lineage bias may emerge through early asymmetries in maternal factor distribution.

In addition to transcriptional divergence, large blastomeres at the 4- and 8-cell stages preferentially exhibited high MAPK14 and RAC1 levels and were highly likely to initiate and maintain cavity formation. These findings conceptually align with previous mouse studies showing that large blastomeres are associated with high CDX2 protein levels, favouring the TE fate^24^. Similarly, blastomeres with large nuclear volumes are also highly likely to contribute to the TE lineage^25^, suggesting that size-dependent cell fate bias is a conserved feature of mammalian embryos. Our results extend these observations to bovine embryos, suggesting that size-dependent molecular and functional asymmetries are conserved across mammals. Importantly, transcriptional variability was not restricted to TE-associated genes in this study. We also detected inter-blastomere differences in the expression of PRDM14, a key regulator of pluripotency and ICM identity^20^. In mouse embryos, PRDM14 is heterogeneously expressed as early as the 4-cell stage, where it collaborates with CARM1-mediated H3R26me2 to promote pluripotency^26^. Although the variability in PRDM14 expression was less pronounced than that in TE marker expression, its presence suggests that early transcriptional asymmetries not only bias TE specification but also influence the emergence of Epi lineages. This supports the notion that early blastomere heterogeneity equally contributes to both embryonic and extraembryonic lineages.

Interestingly, despite the emergence of early molecular and physical asymmetries, developmental plasticity was not immediately lost. Structures derived from the largest blastomeres contained both CDX2-positive trophectodermal cells and SOX2-positive epiblast-like cells, suggesting that the blastomeres biased toward TE-like behaviour contribute to both the embryonic and extraembryonic lineages. This supports the concept of “mosaic plasticity,” wherein early asymmetries bias developmental trajectories without deterministically fixing the cell fate^8, 11^.

The presence of early transcriptional and functional asymmetries in bovine embryos, which exhibit developmental dynamics more comparable to those of humans than to those of rodents, suggests that symmetry-breaking processes are broadly conserved across mammals.

Our findings refine the classical model of early development, which posits uniform totipotency among blastomeres until compaction, highlighting the importance of considering intrinsic heterogeneity during early stage embryo evaluation. These findings have potential implications for assisted reproductive technologies in humans and livestock, as interventions at the cleavage stage may inadvertently disrupt the emerging lineage bias^27–29^.

In summary, transcriptional differences between sister blastomeres emerged progressively before compaction in bovine embryos, similar to that observed in rodent models. These early transcriptional asymmetries possibly bias lineage specification, suggesting the existence of a conserved mechanism underlying early cell-fate decisions across mammals. By extending these insights to a non-rodent model developmentally close to humans, this study provides a valuable framework for advancing basic developmental biology and assisted reproductive technologies.

## Methods

### Reagents

All reagents were purchased from Sigma-Aldrich (St. Louis, MO, USA), unless otherwise stated.

### Oocyte collection

Ovaries were obtained from the Japanese Black or crossbred (Holstein × Japanese Black) cows at a local abattoir and transported to the laboratory at the Tokyo University of Agriculture and Technology. Upon arrival, the ovaries were rinsed with and maintained in 0.9% saline solution (Nippon Zenyaku Kogyo, Fukushima, Japan) at 38.5°C. Cumulus–oocyte complexes (COCs) were aspirated from 2–6-mm antral follicles using a 19G needle attached to a 10-mL syringe.

### In vitro maturation

In vitro maturation of COCs was performed using TCM-199 with 25 mM HEPES (Gibco, Thermo Fisher Scientific, MA, USA) supplemented with 5% calf serum (CS; Gibco) and 0.1LIU/mL recombinant human follicle-stimulating hormone (Follistim; MSD, Tokyo, Japan)^30^. After washing twice with the maturation medium, 50 COCs were cultured in 500LµL of the same medium in 4-well culture plates (Thermo Fisher Scientific) and overlaid with mineral oil (FUJIFILM Wako Pure Chemical Corporation, Osaka, Japan) at 38.5□°C in a humidified atmosphere of 5% CO2 in air for 22□h.

### In vitro fertilization

*In vitro* fertilization was performed as previously described^30^. Frozen semen of a Japanese Black bull was thawed at 37□°C in a water bath and layered into 3LmL of 90% Percoll solution, followed by centrifugation at 740□×□*g* for 10 min. The pellet was resuspended in the Brackett–Oliphant (BO) medium^31^ containing 20□mM hypotaurine, 10□µg/mL heparin (heparin sodium injection; 5,000 U/5□mL; Mochida Pharmaceutical, Tokyo, Japan), and 20 mg/mL bovine serum albumin (BSA; crystallised and lyophilised) and washed via centrifugation at 540□×□*g* for 5□min. The final sperm concentration was adjusted to 3□×□10□/mL in the BO medium supplemented with 20Lmg/mL BSA. In vitro fertilization was performed in 100-µL droplets covered with mineral oil in 35-mm culture dishes (Nunc), each containing 20 COCs. After washing twice with the BO medium containing 10□mg/mL BSA, the COCs were co-incubated with sperms at 38.5□°C in 5% CO_2_ in air for 6□h.

### In vitro culture

After fertilisation, the COCs were denuded of the remaining cumulus cells and sperms via gentle pipetting in the CR1aa medium supplemented with amino acids and 5% CS. Embryos were cultured in groups of 50 in 500□µL of the same medium in 4-well plates and overlaid with mineral oil. In vitro culture was performed at 38.5□°C under low oxygen conditions (5% O_2_, 5% CO_2_, and 90% N_2_).

### Blastomere isolation

Blastomeres were separated from the 2-, 4-, and 8-cell stage embryos 28, 36, and 48Lhpi, respectively^32^. Only the 2-cell stage embryos at 28 hpi were used for subsequent 4- and 8-cell stage embryo collection to avoid abnormal cleavage-stage embryos. Zona pellucida was removed via treatment with 0.25% pronase (actinase E; Kaken Pharmaceutical, Tokyo, Japan) dissolved in phosphate-buffered saline (Gibco). The embryos were further transferred to the CR1aa medium containing 10% CS and mechanically dissociated into individual blastomeres via gentle pipetting using a glass capillary.

### Single-blastomere RNA-sequencing

For transcriptomic analysis, 90 samples, including 10 zygotes, 20 blastomeres from 10 2-cell embryos, 20 blastomeres from five 4-cell embryos, and 40 blastomeres from five 8-cell embryos, were collected. Blastomeres were isolated as described above. Before sampling, each blastomere was imaged to record its diameter using the ImageJ software (NIH), and sister blastomeres from the same embryo were tracked for within-embryo comparisons.

The overall protocol was adapted from our previously described method for single-cell analysis^33^, with some modifications. Total RNA extraction, reverse transcription, and cDNA library preparation were performed using the SMART-Seq HT PLUS Kit (Takara Bio, Shiga, Japan). cDNA quality was assessed using the Agilent 2200 TapeStation system with the High Sensitivity D5000 ScreenTape (Agilent Technologies, CA, USA). Sequencing was conducted on the NovaSeq 6000 platform (Illumina, CA, USA) using 150□bp paired-end reads.

Adapter trimming was performed using Trim Galore, and sequence quality was assessed using FastQC. The reads were aligned to the bovine reference genome (ARS-UCD1.2/bosTau9) using STAR, and transcript quantification was performed using RSEM. PCA was conducted using the prcomp function and hierarchical clustering was performed using the pvclust package in R.

### Assessment of transcriptional asymmetry

To assess the transcriptional asymmetry within embryos, gene-level variability was quantified as SS and partitioned into between-embryo and within-embryo (SSwe) components, as previously described^10^. Genes with the highest SSwe/SS ratios (≥ 0.99 for the 2-cell stage and ≥ 0.70 for the 4- and 8-cell stages) were subjected to KEGG pathway enrichment analysis. These thresholds were empirically determined based on the distribution of the SSwe/SS values at each developmental stage (Figure L2a).

At the 8-cell stage, blastomeres were further classified into high and low CDX2-expressing groups using a threshold of transcripts per million (TPM) ≥ 2.0. PCA and KEGG analyses were conducted using iDEP 0.96^34^.

### Cavity formation assessment

To evaluate the relationship between blastomere size and developmental potential, we assessed cavity formation in the blastomeres isolated from 8-cell embryos (see above for imaging and measurement details in Fig.4a). Each blastomere was individually cultured in a microwell of the LinKID micro25 dish (Dai Nippon Printing, Tokyo, Japan) containing 125 µL of the CR1aa medium supplemented with 10% CS and overlaid with mineral oil (FUJIFILM Wako Pure Chemical Corporation).

Time-lapse monitoring was performed using a real-time embryo culture observation system (CCM-MULTI; Astec, Fukuoka, Japan), with images captured every 15 min using a 10× objective lens^30^. Embryos were monitored up to 192 hours post-insemination (hpi), and classified based on whether they had formed a cavity at any point during this period. The data were further used to examine whether blastomere size is associated with the ability to initiate cavity formation.

## Immunofluorescence staining

Immunofluorescence staining for CDX2 or SOX2 was performed on embryos derived from single 8-cell-stage blastomeres that had formed a cavity at 192 hpi, to assess their differentiation into the trophectoderm (TE) and inner cell mass (ICM), respectively. Embryos were washed 0.2% PVA-PBS and fixed in 4% paraformaldehyde (PFA)-PBS at room temperature for 60 minutes. Fixed embryos were transferred into 0.2% PVA-PBS and stored at 4°C for up to one week. Fixed or stored embryos were permeabilized in 0.2% Triton X-100-PBS at room temperature for 60 minutes, followed by three 10-minute washes in a washing buffer containing 0.1% Triton X-100 and 0.3% bovine serum albumin (BSA)-PBS at room temperature. Blocking was then performed for 45 minutes at room temperature using Blocking One (Nacalai Tesque, Kyoto, Japan) diluted fivefold in PBST (0.05% Tween 20-PBS). Embryos were incubated overnight at 37°C with rabbit monoclonal anti-CDX2 antibody (ab76541; Abcam, Cambridge, UK) diluted 1:300, or at 4°C with rabbit monoclonal anti-SOX2 antibody (ab92494; Abcam) diluted 1:1000. All primary antibodies were diluted in Blocking One diluted 20-fold in PBST. After five 10-minute washes at room temperature with the washing buffer, embryos were incubated with Alexa Fluor™ 555 goat anti-rabbit IgG (A21428; Invitrogen, Thermo Fisher Scientific, MA, USA) diluted 1:400 for 30 minutes at room temperature. From this step onward, all procedures were conducted under light-protected conditions. Following five additional 10-minute washes at room temperature, nuclear staining was performed using Hoechst 33342 (25 μg/mL in 0.2% PVA-PBS) at room temperature for 5 minutes. Finally, embryos were washed three times in 0.2% PVA-PBS and mounted onto glass slides using VECTASHIELD® Mounting Medium (Vector Laboratories, CA, USA), with a coverslip supported by eight pillars made from a mixture of Vaseline (FUJIFILM Wako Pure Chemical Corporation) and liquid paraffin. The edges of the coverslip were sealed with nail polish. Observations were conducted using a confocal laser scanning microscope (LSM710; Carl Zeiss, Baden-Württemberg, Germany) or a fluorescence phase-contrast microscope (BZ-9000; Keyence, Osaka, Japan), and cell counting was performed using Imaris software (Carl Zeiss).

### Statistical analysis

Gene expression changes between the smallest and largest blastomeres within each embryo at the 4- and 8-cell stages were compared using the linear mixed-effects model. “Group” (smallest or largest) was treated as a fixed effect, and “embryo ID” was considered as a random effect to account for within-embryo pairing. The analysis was conducted in Python (version 3.12.3) using the statsmodels package. Cavity formation rates were compared using Fisher’s exact test implemented in R (version 4.3.1) with the fisher.test function from the base stats package. Statistical significance was set at P < 0.05.

## Data availability statement

The RNA-seq data generated in this study have been deposited in the Gene Expression Omnibus (GEO) under accession number GSE301333 and remain private until formal publication.

## Acknowledgements

This work was supported by JSPS KAKENHI Grant Number JP23K23760 to S.S., and the JRA Livestock Industry Promotion Project to S.S. We would like to thank Editage (www.editage.com) for English language editing.

## Author contributions

S.S. and H.K. conceptualized the study. H.K. conducted the experiments. S.S., H.K., and D.M. conducted data analysis. M.K. and A.K. provided support for RNA-seq analysis. S.S., H.K., and D.M wrote the original draft of the manuscript. S.S., H.K., and D.M. reviewed and edited the manuscript. S.S. supervised the project and secured funding.

## References

1. Tarkowski, A.K. Experiments on the Development of Isolated Blastomeres of Mouse Eggs. Nature 184, 1286–1287 (1959).

2. Rossant, J. & Tam, P.P. Blastocyst lineage formation, early embryonic asymmetries and axis patterning in the mouse. Development 136, 701–713 (2009).

3. Johnson, M.H. & McConnell, J.M. Lineage allocation and cell polarity during mouse embryogenesis. Semin Cell Dev Biol 15, 583–597 (2004).

4. Zernicka-Goetz, M., Morris, S.A. & Bruce, A.W. Making a firm decision: multifaceted regulation of cell fate in the early mouse embryo. Nat Rev Genet 10, 467–477 (2009).

5. Morris, S.A., Guo, Y. & Zernicka-Goetz, M. Developmental plasticity is bound by pluripotency and the Fgf and Wnt signaling pathways. Cell Rep 2, 756–765 (2012).

6. Plusa, B. et al. The first cleavage of the mouse zygote predicts the blastocyst axis. Nature 434, 391–395 (2005).

7. Bischoff, M., Parfitt, D.E. & Zernicka-Goetz, M. Formation of the embryonic-abembryonic axis of the mouse blastocyst: relationships between orientation of early cleavage divisions and pattern of symmetric/asymmetric divisions. Development 135, 953–962 (2008).

8. Goolam, M. et al. Heterogeneity in Oct4 and Sox2 Targets Biases Cell Fate in 4-Cell Mouse Embryos. Cell 165, 61–74 (2016).

9. White, M.D. et al. Long-Lived Binding of Sox2 to DNA Predicts Cell Fate in the Four-Cell Mouse Embryo. Cell 165, 75–87 (2016).

10. Biase, F.H., Cao, X. & Zhong, S. Cell fate inclination within 2-cell and 4-cell mouse embryos revealed by single-cell RNA sequencing. Genome Res 24, 1787–1796 (2014).

11. Shi, J. et al. Dynamic transcriptional symmetry-breaking in pre-implantation mammalian embryo development revealed by single-cell RNA-seq. Development 142, 3468–3477 (2015).

12. Casser, E., Israel, S., Schlatt, S., Nordhoff, V. & Boiani, M. Retrospective analysis: reproducibility of interblastomere differences of mRNA expression in 2-cell stage mouse embryos is remarkably poor due to combinatorial mechanisms of blastomere diversification. Mol Hum Reprod 24, 388–400 (2018).

13. Memili, E. & First, N.L. Zygotic and embryonic gene expression in cow: a review of timing and mechanisms of early gene expression as compared with other species. Zygote 8, 87–96 (2000).

14. Telford, N.A., Watson, A.J. & Schultz, G.A. Transition from maternal to embryonic control in early mammalian development: a comparison of several species. Mol Reprod Dev 26, 90–100 (1990).

15. Suzuki, R., Okada, M., Nagai, H., Kobayashi, J. & Sugimura, S. Morphokinetic analysis of pronuclei using time-lapse cinematography in bovine zygotes. Theriogenology 166, 55–63 (2021).

16. Berg, D.K. et al. Trophectoderm lineage determination in cattle. Dev Cell 20, 244–255 (2011).

17. Niakan, K.K., Han, J., Pedersen, R.A., Simon, C. & Pera, R.A. Human pre-implantation embryo development. Development 139, 829–841 (2012).

18. Negrón-Pérez, V.M., Zhang, Y. & Hansen, P.J. Single-cell gene expression of the bovine blastocyst. Reproduction 154, 627–644 (2017).

19. Lu, C.W. et al. Ras-MAPK signaling promotes trophectoderm formation from embryonic stem cells and mouse embryos. Nat Genet 40, 921–926 (2008).

20. Yamaji, M. et al. PRDM14 ensures naive pluripotency through dual regulation of signaling and epigenetic pathways in mouse embryonic stem cells. Cell Stem Cell 12, 368–382 (2013).

21. Piotrowska-Nitsche, K., Perea-Gomez, A., Haraguchi, S. & Zernicka-Goetz, M. Four-cell stage mouse blastomeres have different developmental properties. Development 132, 479–490 (2005).

22. Wang, J. et al. Asymmetric Expression of LincGET Biases Cell Fate in Two-Cell Mouse Embryos. Cell 175, 1887–1901.e1818 (2018).

23. Graf, A. et al. Fine mapping of genome activation in bovine embryos by RNA sequencing. Proc Natl Acad Sci U S A 111, 4139–4144 (2014).

24. Dietrich, J.E. & Hiiragi, T. Stochastic patterning in the mouse pre-implantation embryo. Development 134, 4219–4231 (2007).

25. Nunley, H. et al. Nuclear instance segmentation and tracking for preimplantation mouse embryos. Development 151 (2024).

26. Burton, A. et al. Single-cell profiling of epigenetic modifiers identifies PRDM14 as an inducer of cell fate in the mammalian embryo. Cell Rep 5, 687–701 (2013).

27. Kirkegaard, K., Hindkjaer, J.J. & Ingerslev, H.J. Human embryonic development after blastomere removal: a time-lapse analysis. Hum Reprod 27, 97–105 (2012).

28. Kalma, Y. et al. Optimal timing for blastomere biopsy of 8-cell embryos for preimplantation genetic diagnosis. Hum Reprod 33, 32–38 (2018).

29. Park, J.H. et al. Rapid sexing of preimplantation bovine embryo using consecutive and multiplex polymerase chain reaction (PCR) with biopsied single blastomere. Theriogenology 55, 1843–1853 (2001).

30. Sugimura, S. et al. Time-lapse cinematography-compatible polystyrene-based microwell culture system: a novel tool for tracking the development of individual bovine embryos. Biol Reprod 83, 970–978 (2010).

31. Brackett, B.G. & Oliphant, G. Capacitation of rabbit spermatozoa in vitro. Biol Reprod 12, 260–274 (1975).

32. Sugimura, S. et al. Effect of embryo density on in vitro development and gene expression in bovine in vitro-fertilized embryos cultured in a microwell system. J Reprod Dev 59, 115–122 (2013).

33. Tsuji, H. et al. Compatibility of dry incubator on in vitro production of bovine embryos. Theriogenology 232, 117–123 (2025).

34. Ge, S.X., Son, E.W. & Yao, R. iDEP: an integrated web application for differential expression and pathway analysis of RNA-Seq data. BMC Bioinformatics 19, 534 (2018).

