## Supplementary figures and images for "Early transcriptional divergence underlies cell fate bias in bovine embryos"

### Supplemental Fig. 1

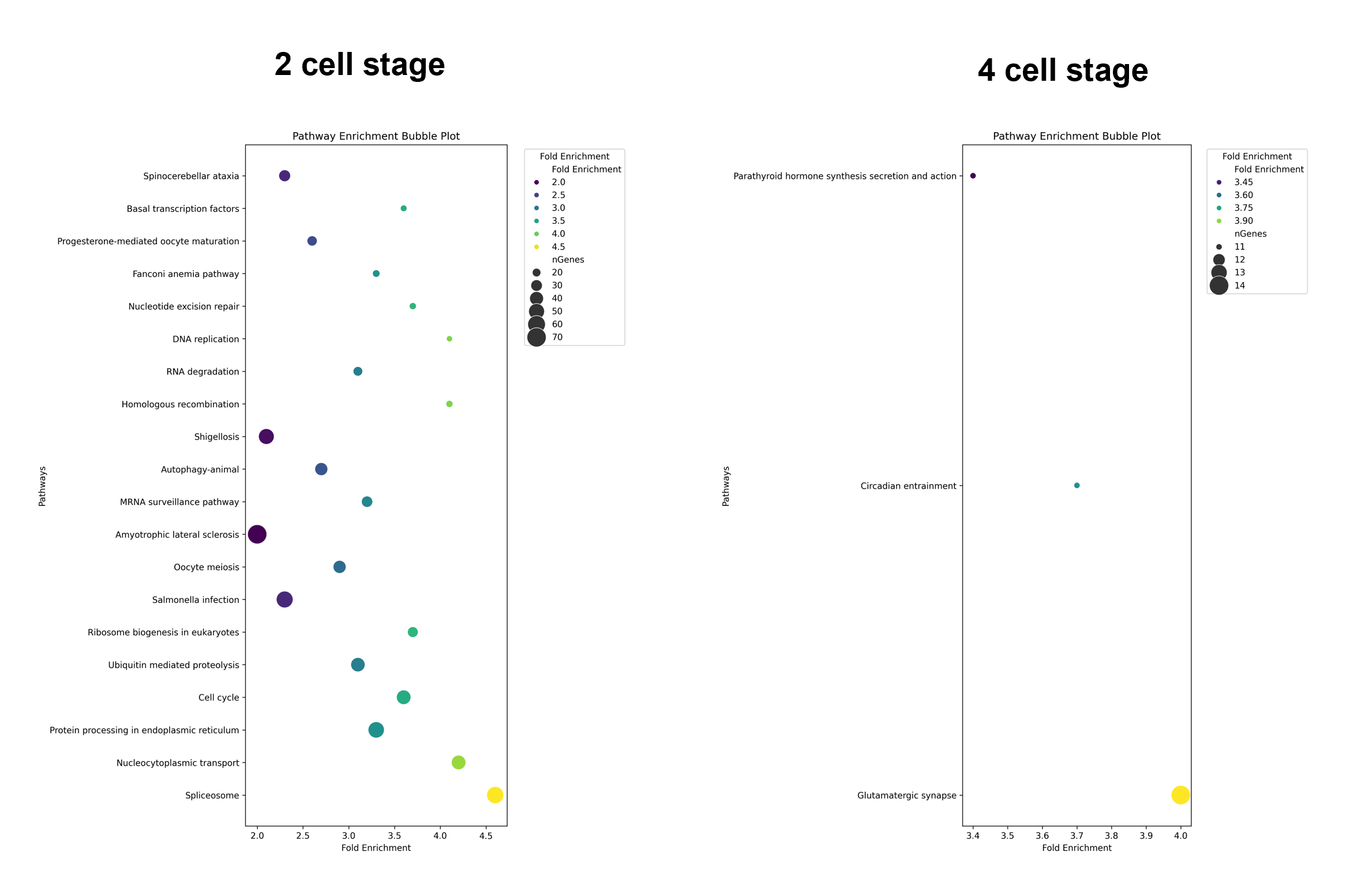
