## Supplemental Table 1 for "Early transcriptional divergence underlies cell fate bias in bovine embryos"

Supplementary Table 3. Differentially expressed genes (DEGs) between the smallest and largest blastomeres at the 4-cell and 8-cell stages.
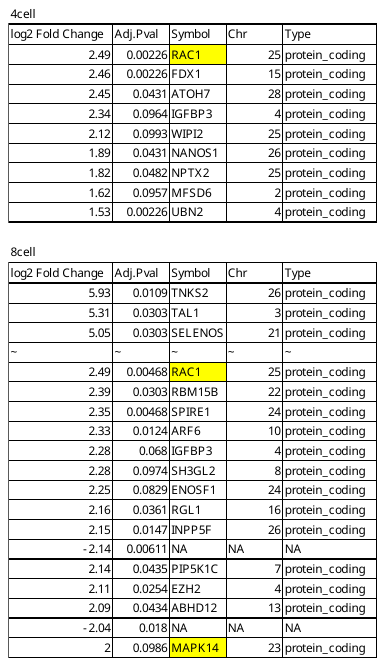
